# Genome-guided discovery of natural products through multiplexed low coverage whole-genome sequencing of soil Actinomycetes on Oxford Nanopore Flongle

**DOI:** 10.1101/2021.08.11.456034

**Authors:** Rahim Rajwani, Shannon I. Ohlemacher, Gengxiang Zhao, Hong-bing Liu, Carole A. Bewley

## Abstract

Genome-mining is an important tool for discovery of new natural products; however, the number of publicly available genomes for natural product-rich microbes such as Actinomycetes, relative to human pathogens with smaller genomes, is small. To obtain contiguous DNA assemblies and identify large (ca. 10 to greater than 100 Kb) biosynthetic gene clusters (BGCs) with high-GC (>70%) and -repeat content, it is necessary to use long-read sequencing methods when sequencing Actinomycete genomes. One of the hurdles to long-read sequencing is the higher cost.

In the current study, we assessed Flongle, a recently launched platform by Oxford Nanopore Technologies, as a low-cost DNA sequencing option to obtain contiguous DNA assemblies and analyze BGCs. To make the workflow more cost-effective, we multiplexed up to four samples in a single Flongle sequencing experiment while expecting low-sequencing coverage per sample. We hypothesized that contiguous DNA assemblies might enable analysis of BGCs even at low sequencing depth. To assess the value of these assemblies, we collected high-resolution mass-spectrometry data and conducted a multi-omics analysis to connect BGCs to secondary metabolites.

In total, we assembled genomes for 20 distinct strains across seven sequencing experiments. In each experiment, 50% of the bases were in reads longer than 10 Kb, which facilitated the assembly of reads into contigs with an average N50 value of 3.5 Mb. The programs antiSMASH and PRISM predicted 629 and 295 BGCs, respectively. We connected BGCs to metabolites for *N*,*N*-dimethyl cyclic-ditryptophan, a novel lassopeptide and three known Actinomycete-associated siderophores, namely mirubactin, heterobactin and salinichelin.

**Importance:** Short-read sequencing of GC-rich genomes such as Actinomycetes results in a fragmented genome assembly and truncated biosynthetic gene clusters (often 10 to >100 Kb long), which hinders our ability to understand the biosynthetic potential of a given strain and predict the molecules that can be produced. The current study demonstrates that contiguous DNA assemblies, suitable for analysis of BGCs, can be obtained through low-coverage, multiplexed sequencing on Flongle, which provides a new low-cost workflow ($30-40 per strain) for sequencing Actinomycete strain libraries.

## Introduction

Clinical pathogens are increasingly becoming resistant to currently used antimicrobials causing over 700,000 deaths worldwide (1). New antimicrobials are urgently needed to alleviate antimicrobial resistance and prevent deaths per year to rise over 10 million by 2050 (1). One of the prolific sources of new antimicrobials is a group of gram-positive mycelia forming bacteria, the Actinomycetes. Several currently used antibiotics, including vancomycin, rifamycin, and streptomycin are isolated from Actinomycetes and they still hold enormous potential for the future discovery of new medicines (2).

Genome sequencing is now an important component of natural products research. Whole-genome sequencing (WGS) enables identification of the genes responsible for the biosynthesis of natural products (3). Often genes required for the biosynthesis of a natural product positionally cluster on the genome and are referred to as biosynthetic gene clusters (BGCs) (4). The BGC sequences can be used to predict possible structures of the resulting natural product (5), assess novelty of the compound (6) and dereplicate compounds from a strain collection (7). Despite the merits offered by WGS, the number of Actinomycete genomes remains limited. Several rare genera are not represented by a complete genome, and the majority of currently available genomes are sequenced using Illumina short-read technology that results in highly fragmented assemblies. BGCs span multiple contigs in fragmented genome assemblies and cannot be detected or analyzed by commonly used BGC prediction tools such as antiSMASH (8, 9).

Long-read sequencing technologies (e.g. PacBio or Oxford Nanopore Technologies, ONT) produce contiguous genome sequences needed to analyze secondary metabolite gene clusters. Notably, PacBio assemblies achieve consensus accuracy over 99.999%; however, it is generally less accessible due to the upfront cost of sequencing instruments and higher per sample sequencing costs. By contrast, ONT does not require an upfront cost of an expensive sequencing instrument and the devices are inexpensive. Nevertheless, ONT data results in a lower consensus accuracy (99.9%) and often requires polishing with Illumina reads to obtain reference-quality genomes. We hypothesized that while BGC identification requires a contiguous DNA sequence, it might be less affected by the lower consensus accuracy of a Nanopore assembly since most BGC analysis steps involve inferring homology between distantly related amino acid sequences using profile Hidden Markov models. If this is true, contiguous DNA assemblies can be obtained at ca. 10× coverage using ONT, allowing complete genome sequencing at a significantly lower cost. While such ONT sequenced genomes would still require error correction with Illumina reads, they could be used on their own to sequence a strain collection, build a catalog and compare BGCs for dereplication or identification of potentially new compounds, which might be particularly useful to natural product research and drug discovery programs.

To assess the feasibility of obtaining contiguous assemblies from ca. 10× sequencing depth, predicting BGCs, and connecting BGCs to metabolites, we conducted the current multi-omics study. We sequenced 20 new soil-derived Actinomycete strains and analyzed their metabolome using high-resolution mass spectrometry (HRMS). For sequencing, we specifically selected Flongle, a recently launched ONT sequencing device that costs $90 USD and can generate up to 1-2 Gigabases of sequence output. With a typical Actinomycetes genome being 8-10 Mb, a single Flongle experiment might be sufficient to sequence 3-4 strains at 20-30× coverage. Sequencing workflows based on Flongle could be broadly applicable to small and large studies due to the modular experimental design. In the current study, we obtained 300-850 Mb of data per experiment across ten sequencing experiments with read-length N50 values over 10 Kb. Assembling of reads resulted in contiguous assemblies (average contig N50 value = 3.5 Mb and average number of contigs = 47.3). AntiSMASH5 predicted a total of 629 BGCs from these assemblies. Through a combined analysis with metabolomics data, we were able to connect BGCs to their secondary metabolites. The study demonstrates the utility of low coverage nanopore-only assemblies as a rapid and low-cost sequencing option to advance natural product research.

## Results

### An in silico analysis to study the effect of sequencing coverage and read length on BGC detection

We first analyzed what level of sequencing coverage would be sufficient for contiguous assemblies and BGC detection using Oxford Nanopore sequencing. For this purpose, three Actinomycete genomes, previously sequenced at high coverage, were downloaded from the European Nucleotide Archive and their reads were down sampled to 60×, 30×, 15× and 7× coverage (assuming a genome size of 8 Mb) before assembling and detecting BGCs. While actual genome sizes of the three genomes differed (Table S 1), an assumption of a fixed expected genome size of 8 Mb allowed us to determine the utility of a prospective sequencing experiment where Actinomycete genome sizes would not be known. In the down sampling analysis, assembly size and number of predicted genes nearly plateau at ca. 15× coverage. Similarly, a sharp decline in the number of contigs and number of mismatches per 100 Kb was observed at ca. 15-20× coverage (Figure 1). At approximately the same coverage of 15-20×, 72-96% of BGCs were detected by antiSMASH and a further increase in coverage led to detection of only 1-6 additional BGCs (Figure 1). Moreover, most of the BGCs were not located at the edge of a contig, also referred to as complete. Relative to antiSMASH, PRISM predicted a lower number of BGCs. This could be because antiSMASH was run in a ‘relaxed’ mode in this study whereas PRISM does not have this option. Nevertheless, the trend relative to coverage was similar between antiSMASH and PRISM.

**Figure 1:**
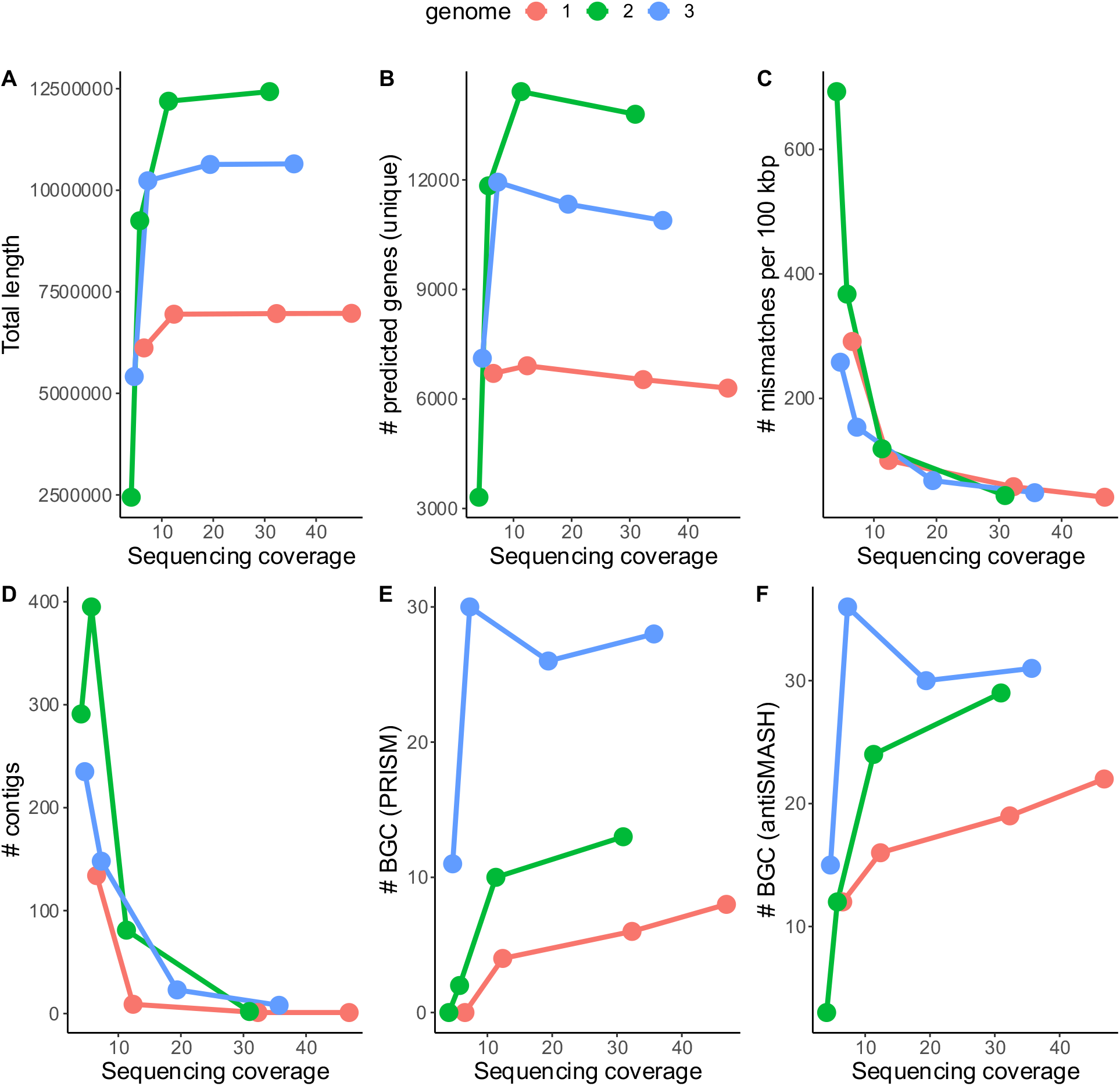
An in silico analysis of the effect of sequencing coverage on assembly quality and BGC prediction. Three genomes previously sequenced at high coverage on ONT MinION/GridION platform were down sampled to the indicated sequencing coverages and then assembled. A-F The quality of the assembly as indicated by total assembly length, number of contigs, number of mismatches per 100 Kb and number of unique predicted genes are shown along with the number of BGCs predicted by antiSmash. The sequencing data for this analysis was downloaded from the European Nucleotide Archive. Accession numbers: 1 = SRR10597857, 2 = SRR9710049, 3 = DRR240480).

Assembly contiguity and therefore BGC detection in a Nanopore sequencing experiment is also related to read length. In another computational experiment, we evaluated whether longer reads might enable a more contiguous DNA assembly and BGC detection at fixed coverage. For this purpose, simulated Nanopore reads of average length 500, 1000, 2000 and 4000 were generated at 10× coverage of a Streptomyces genome (GB4-14) using BadRead. The resulting reads were assembled and analyzed for assembly contiguity and BGC detection. It was observed that an ca. 2-fold increase in average read length was associated with a 2-fold reduction in the number of contigs (Figure S 3). Improved assembly contiguity also led to a reduction in the number of BGCs on contig edges (incomplete) and an increased number of complete BGCs with little to no change of sequencing coverage (Figure S 4).

Overall, these computational experiments suggested contiguous DNA assemblies and complete BGCs can be detected at low sequencing coverage using long reads from Oxford Nanopore Technologies; this should allow for a dramatic reduction in cost per genome through multiplexing. The computational experiments were followed up with prospective sequencing of Actinomycetes genomes using Flongle and more detailed analyses of BGCs described in the following sections.

### Nanopore sequencing, genome assembly and quality assessment

A total of ten sequencing experiments were conducted— each with an attempt to sequence four Actinomycetes strains (Figure 2). Impurities in the starting genomic DNA (as measured by a ratio of the UV absorbance at 260 and 280nm) and low pore occupancy (caused by insufficient loading of the library or inhibition of adapter ligation) resulted in three unsuccessful experiments with a total output <100 megabases (Mb) per experiment. The remaining seven experiments yielded 288-797 Mb over 18-24 hours. The longest read for each sample was over 80 Kb.

**Figure 2:**
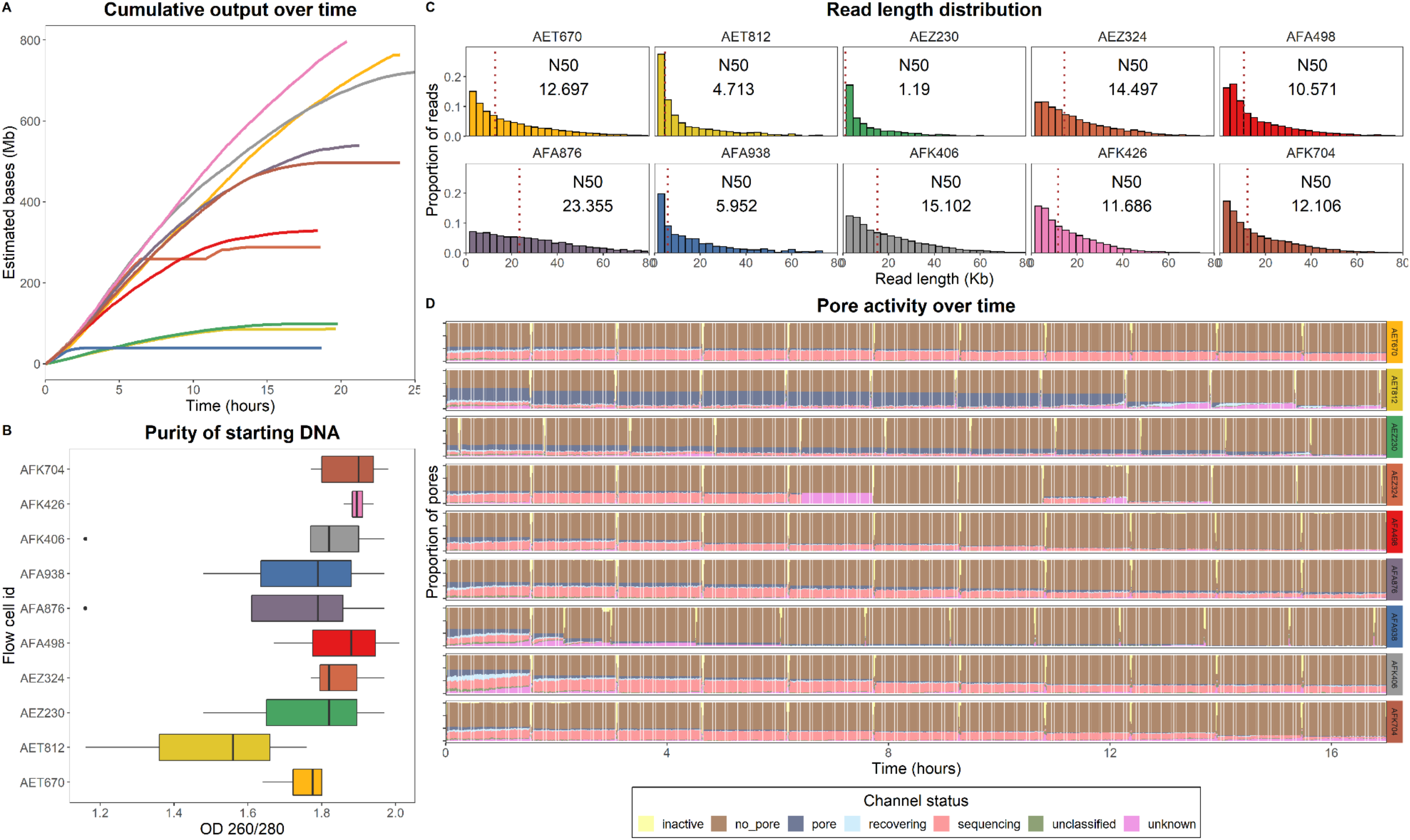
Library preparation and sequencing quality metrics. The data in each panel (A-D) are colored and grouped by flow cell as indicated in panel B. Panel A indicates cumulative output (estimated megabases) over time (hours) across 10 sequencing experiments. Panel B indicates the purity (nanodrop 260/280 ratio) for samples run in each experiment. Panel C indicates read length distribution across experiments. Read N50 value (50% of bases are in reads longer this value) in each experiment is labeled. Panel D shows run performance at the pore level in each experiment except AFK4226. The AFK4226 experiment was interrupted at the end and the instrument did not generate the pore activity metadata to include in this chart. “Sequencing” indicates that the pore is occupied with DNA and is sequencing. “Pore” indicates an empty pore with no DNA, “no pore” indicates an inactive pore (i.e. unavailable for sequencing).

Across the experiments, we tried different buffers for bead-based purification to apply size selection and increase the read length N50 values from standard protocols (Table S 2) (Figure 2). One of our initial experiments using 0.5× of the standard buffer concentration was not successful resulting in read N50 values of 1-1.6 Kb for three out of five samples sequenced in the experiment. In two subsequent successful experiments (AET670and AFK704), we utilized 0.15× of a modified buffer containing 0.5 M MgCl_2_ + 5% PEG in TE buffer (10 mM Tris-Cl pH 8.0, 1 mM EDTA) for bead-based purification after barcodes ligation as described previously (10). Read length N50 values in these experiments were 11.6-15.1 Kb (Figure 2). In three experiments (AEZ324, AFA498 and AFA876), the concentration of the modified buffer-based size selection was reduced to 0.1× which led to a further increase in read length N50s (10.5-23.3 Kb) accompanied by an increased sample loss. Application of size selection after barcodes ligation ensures approximately equal fragment lengths for pooling of samples and adapter ligation. However, ligation of barcodes could be less efficient to longer fragments if shorter fragments are present in the mixture. We tested buffer-based size selection (0.1× beads in modified buffer) before barcode ligation in later experiments (flow cell ids AFK426 and AFK406) (Table S 2). A more consistent output was observed, possibly due to more efficient barcode/adapter ligation to longer DNA fragments.

Across the seven successful runs, 3,814,434,062 bases in 751,459 reads were generated. Upon demultiplexing, the median number of bases per sample was 77.5 Mb (theoretical coverage of 9.5× with an expected 8 Mb genome size). Three strains were sequenced at <2.5× theoretical coverage (<20Mb per strain) and were excluded from further analysis. Subsequently, 25 samples (20 distinct isolates) were de novo assembled with Canu and polished with Racon and medaka (Figure 3). The median length of the obtained assemblies was 8.5 Mb (average: 7.9 Mb, maximum: 9.4 Mb), typical of Actinomycete genome size. The only exception was a 3 Mb assembly for GB8-002 which was also sequenced at the least coverage (4.0×) (Figure 3).

**Figure 3:**
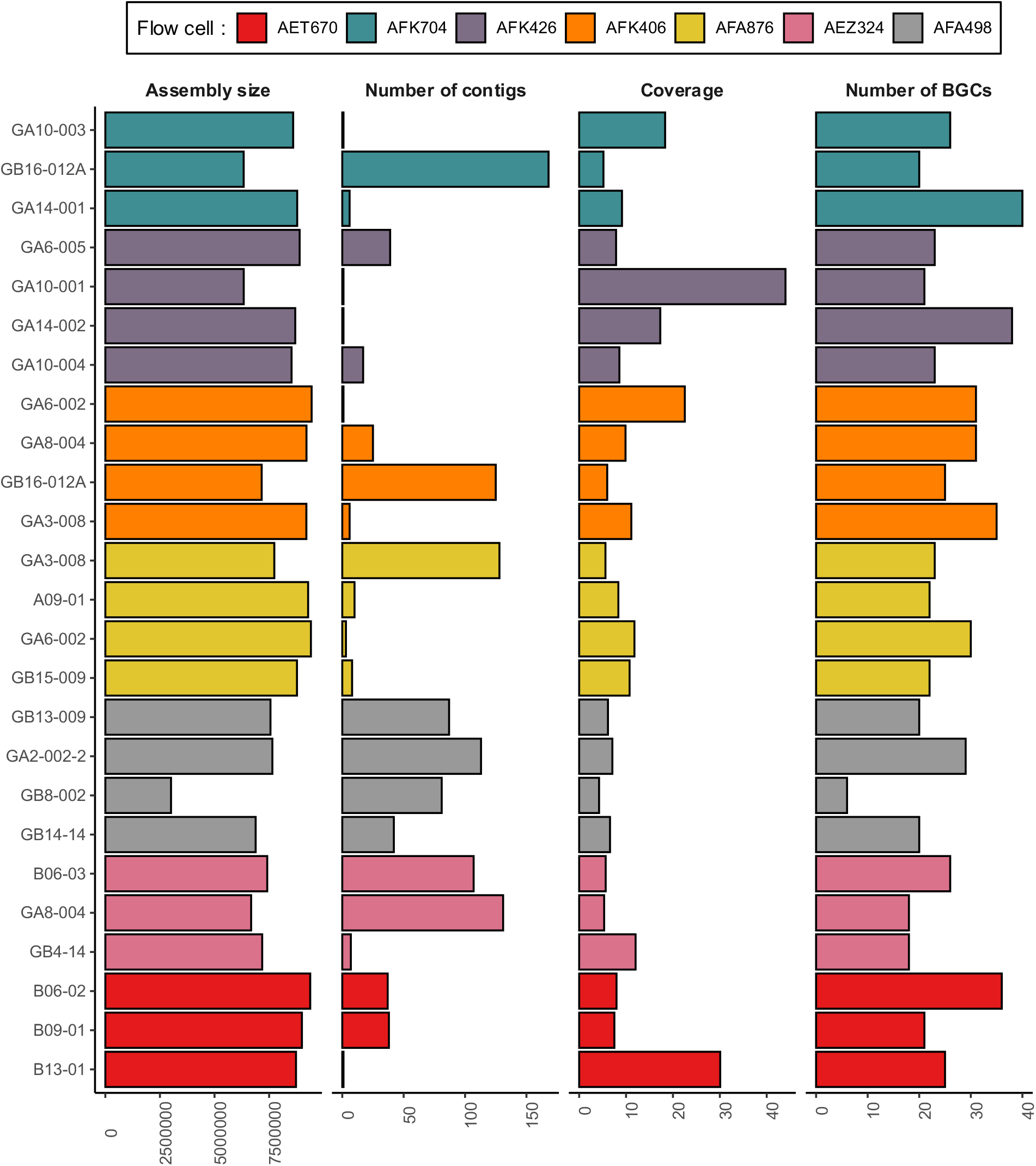
Characteristics of the genome assemblies obtained through low-pass multiplexed sequencing on Flongle. Each bar represents a sample. Bars are grouped and colored by flow cells (individual sequencing experiment).

We assessed the accuracy and quality of these low coverage genomes by comparing them with genomes sequenced at high coverage on MinION or PacBio. In particular, two strains, GA3-008 and GB4-14 were previously sequenced by our lab at 10-fold higher coverage using MinION and PacBio, respectively (Table 1). Despite the lower sequence coverage, the genomes’ contiguity was only slightly affected on flongle and all were assembled into <10 contigs. The size of the assembly differed by 6.1 Kb (GA3-008) and 19.6 Kb (GB4-14) due to insertion/deletion (indel) errors. Despite many mismatches and indel errors, 87.5-100% of the BGCs detected in the MinION or PacBio assemblies were also detected in these Flongle assemblies by antiSMASH.

**Table 1:** Quality assessment of genomes of two strains (GA3-008 and GB4-14) obtained from low coverage flongle assemblies, compared to PacBio and MinION assemblies.

| **Attribute** | **GA3-008** |  | **GB4-14** |  |
| --- | --- | --- | --- | --- |
|  | Flongle multiplex | Minion singleplex | Flongle multiplex | PacBio |
| **Est. coverage** | 15.3 | 121.65 | 14 | 200 |
| **# contigs** | 6 | 1 | 7 | 2 |
| **Assembly size** | 9205503 | 9199325 | 7183038 | 7163416 |
| **# mismatches\*** | 3000 | - | 19227 | - |
| **# indels \*** | 13803 | - | 14484 | - |
| **# BGCs** | 35 | 40 | 18 | 18 |

\*Number of mismatches and indels in the Flongle multiplexed assemblies are relative to the same strain sequenced by either MinION or PacBio.

### Taxonomy and BGCs

antiSMASH predicted 629 BGCs of 29 different types across all assembled genomes from this study (Figure S 5). Seventy-nine percent (497/629) of these BGCs were complete (i.e. not located on a contig edge). There was a median of 23 BGCs per strain. The number of non-ribosomal peptide synthases (NRPSs) and terpenes detected were 2-3 times greater overall than other BGC types; however, this may be because antiSMASH does not subdivide these BGC types into defined sub-classifications, like it does for the RiPPs (LAP, lanthipeptide, etc.) and PKS (type I, II, etc.) categories. In addition to antiSMASH, 295 BGCs were predicted using PRISM software. A unique feature of PRISM is that it enables chemical structure prediction from BGCs (11). In the current dataset, PRISM generated a predicted chemical structure for 180 out of 295 predicted BGCs.

The taxonomic identification based on 16S rRNA sequences extracted from the WGS revealed that the dataset comprises 11 different Actinomycete species belonging to four genera (Table S 3). It consisted of eight *Amycolatopsis*, nine *Streptomyces*, four *Lentzea* and four *Nocardia* species. Some species were overrepresented in the dataset. *Amycolatopsis lurida* and *Streptomyces tendae* were each represented with four strains and *Lentzea violacea* with two strains (Table S 3). The biosynthetic diversity between strains was high with members of the same species, sharing >99% identity in their 16S sequences, differing by up to 20 BGCs (Figure S 6). The biosynthetic diversity of BGCs within species also varied in some cases. For instance, strains of *Streptomyces tendae*, *Lentzea violacea* or *Streptomyces kanamyceticus* were more diverse within the species than strains of *Amycolatopsis lurida.* Different species of the same genus also encoded 10-15 different BGCs on average.

### Predicting metabolites from BGCs – paired analysis of genome and secondary metabolites

#### Insilco PRISM-predicted chemical structures

The PRISM predicted BGCs detected from low coverage assemblies were analyzed further to determine whether they could be linked to a metabolite. We conducted a paired analysis by collecting MS/MS spectra for extracts from strain cultures grown in ISP1 and R2A media. MS/MS spectra were queried using molDiscovery against a database of PRISM predicted chemical structures from BGCs (12). The database comprised 1,177 structures generated from 180 BGCs sequenced in this study. A total of 18 predicted structures matched to MS/MS spectra collected from the strains at false discovery rate (FDR) < 1% and p-value < e^−10^. Three of these matches were also detected in media blanks and were excluded from the analysis. These three metabolites corresponded to tryptophan, and the dipeptides Pro-Val and Phe-Val.

Two of eighteen matched structures were predicted from a cyclic dipeptide BGC in *Amycolatopsis* strain GA6-002 and were detected in metabolite extracts from the same strain (Figure 4). The two gene BGC encoded a cyclic dipeptide synthase (CDPS) and an *N*-methyl transferase. Both of the amino acyl tRNA binding pockets of the CDPS had a specificity signature for tryptophan tRNA. Based on sequence information, cyclic-di-tryptophan c(WW) was predicted as a possible metabolite that can be methylated on nitrogen by the *N*-methyl transferase as reported previously for *Actinosynnema mirum* (13). In all chemical extracts from GA6-002, a metabolite matching the precursor *m/z* and predicted MS/MS fragmentation pattern for c(WW) was detected. In addition, in the ethyl acetate extracts from ISP1 cultures the *N*-methylated metabolite c(WW)Me2 was also detected.

**Figure 4:**
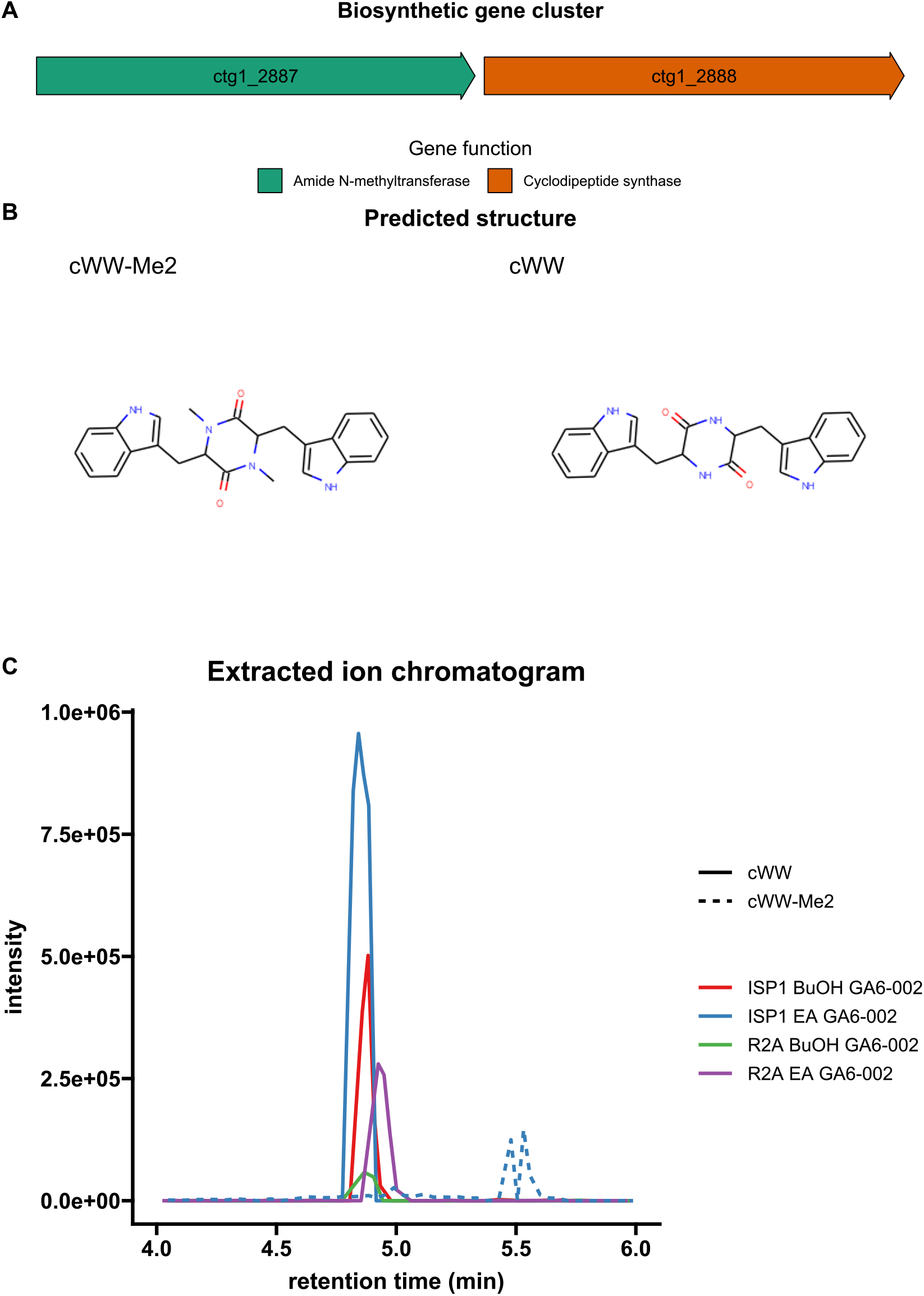
An *N*-methylated cyclo(Trp-Trp) cyclic dipeptide detected in *Amycolatopsis* sp. GA6-002 and its corresponding BGC. A) Cyclic dipeptide BGC and B) the structures predicted by PRISM based on predicted specificity of the cyclic dipeptide synthase. C) Extracted ion chromatogram for cyclic di tryptophan (cWW) and its *N*-methylated derivative (cWW-Me2) observed in four separate crude extracts.

An additional mass spectral match was detected for a small, glycosylated polyketide in GB4-14 consisting of a propionate unit and an actinosamine sugar moiety (Figure S 7), resembling a putative shunt product of a larger polyketide. The corresponding BGC includes additional PKS modules that were not accounted for in the structure predicted by PRISM. More complex structures that better resemble final products of PKS pathways were predicted from a more contiguous Flongle (Figure 3) or PacBio assembly (Table 1) of the same strain but were not detected in the metabolite extracts. The final product of this BGC, predicted from PacBio sequenced genome, was therefore regarded as not detected.

#### RiPPs

We conducted a second analysis to query MS/MS spectra for post-translationally modified precursor peptides from RiPPS using MetaMiner (14). All open reading frames shorter than 600 nt were extracted from 43 antiSmash predicted RiPP BGCs (16 lanthipeptide, 4 LAP, 13 lassopeptides, 10 thiopeptides) and included in this analysis (Figure S 5). We observed a single high confident match for a class-II lassopeptide BGC in the *Amycolatopsis* sp. GA6−002 (Figure 5). The BGC encoded all essential elements for lasso peptide biosynthesis including precursor peptide, asparagine synthetase (SMCOG1177 - essential for macrolactam formation), lassopeptide transglutamase protease (PF13471 - leader peptide cleavage) RiPP recognition element (PF05402), and ABC transporter (SMCOG1288 and SMCOG1000) (15, 16)(Figure 5) [14, 15]. The precursor *m/z* (1041.504 [M+2H]^2+^ and 694.672 [M+2H]^3+^) of the matched spectra was consistent with the predicted core peptide after loss of one water molecule (−18.010). The MS^2^ fragmentation pattern further indicated abundant ions matching *m/z* for y6 and y7 ions. The 16 amino acid core peptide sequence (GYPWWDNRDIFGGRTFL) is a novel lassopeptide variant with 76% amino acid identity to propeptin, an endopeptidase inhibitor (17). The analysis was also repeated for RiPP BGCs predicted by PRISM and no matches were detected with p-value lower than e^−10^.

**Figure 5:**
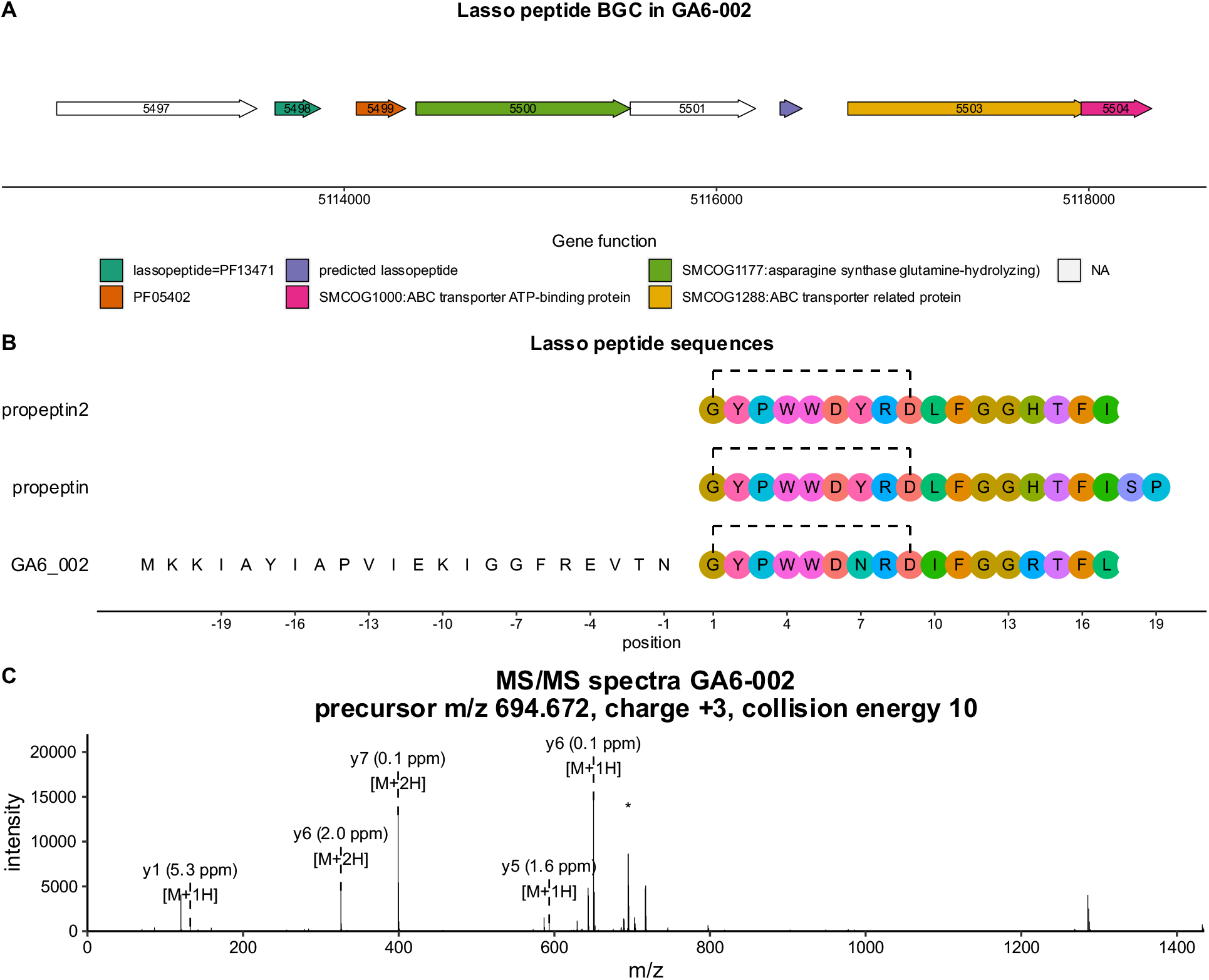
A new lasso peptide variant identified in GA6-002. A) Lasso peptide biosynthetic gene cluster. Genes are colored according to their functions. B) Lassopeptide sequences in propeptin2, propeptin and GA6-002 (this study). Amino acid positions for leader sequence are shown as negative numbers. A BGC for propeptin 2 and propeptin has yet to be characterized and therefore the leader sequences are not shown for them. Dashed line between Gly and Asp shows the position of crosslink. C) MS/MS spectra that matches to the post-translationally modified predicted core sequence in GA6-002.

#### Known metabolites and their BGCs

An important application of genome sequencing is to understand the biosynthesis of known natural products. Similarity to characterized BGCs can also be used for strain dereplication. To assess this application on the current sequencing data, we screened MS/MS spectra for known natural products in the Natural Product Atlas database (3) (29,006 compounds) using molDiscovery (12). Subsequently, we analyzed genome sequences to confirm the presence of the corresponding BGCs. A total 324 significant matches to known compounds were detected (p-value < e^10^). Of these, 30 had a reference MS/MS spectrum available in GNPS. We compared the spectra observed in our dataset to the reference spectra available in GNPS and found highly similar MS/MS spectra (Figure S 8).

Twenty-one of the 324 identified compounds had a previously characterized BGC in the MiBiG database (4). Twelve of these were known Actinomycete natural products; therefore, a higher sequence similarity could be expected. The other nine were compounds isolated from diverse bacterial genera, including those from Gram-negative bacteria and the phylum Cyanobacteria. The presence in our genomes of homologous BGCs for four known Actinomycetes compounds could be confirmed using BLAST sequence similarity searches (Figure 6). These compounds included *N*-acetyl tryptophan and the siderophores heterobactin A, mirubactin and salinichelin. From the BGC comparisons illustrated in Figure 6, it is evident that the nanopore-sequenced genomes from this study can have sequencing artifacts resulting in fragmentation of large genes into multiple small ORFs (see for example the comparison of mirubactin). However, matches to homologous BGCs in MiBiG were easily identified by the high sequence identity between genes, shared functions, and synteny.

**Figure 6:**
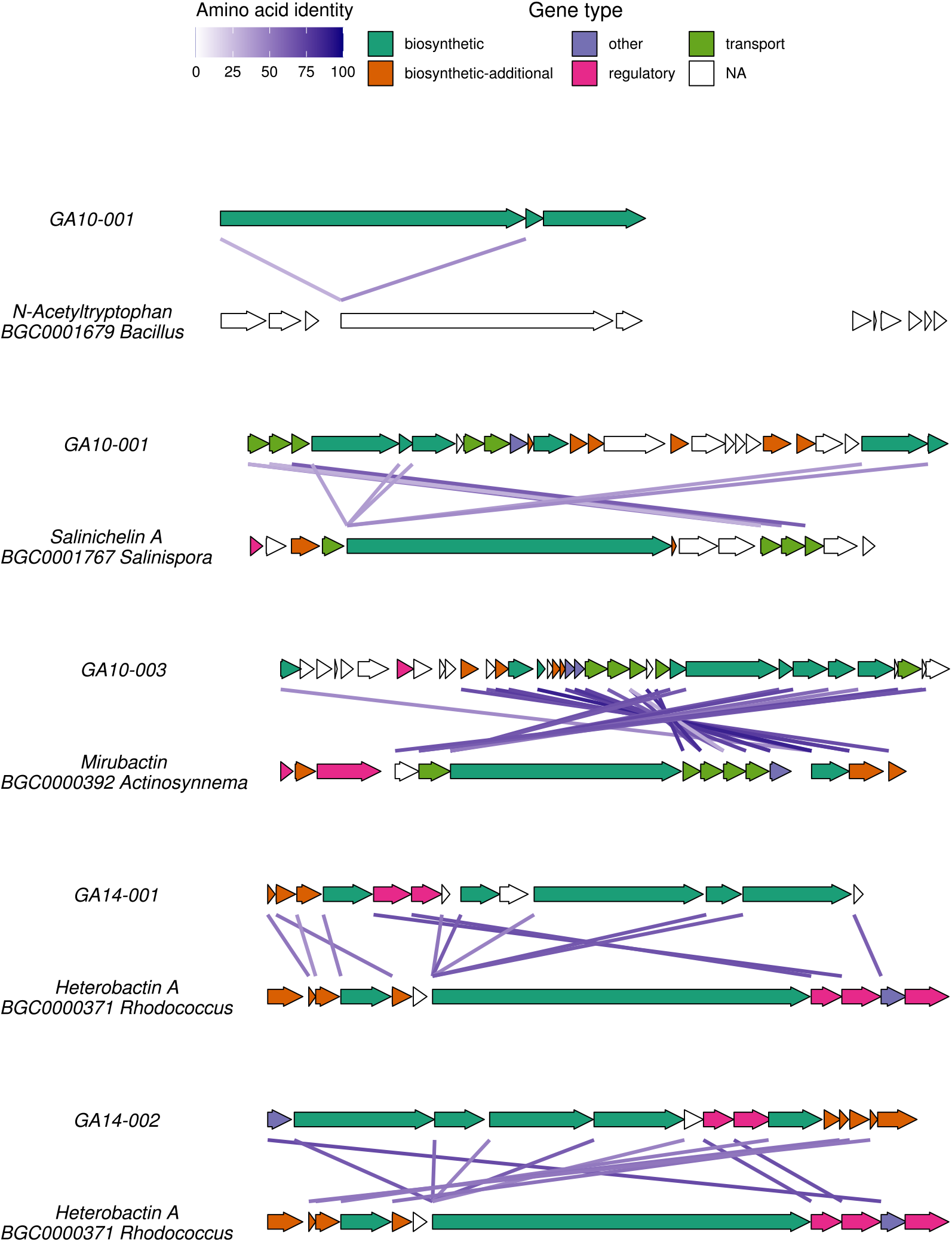
Homologous BGCs mapped for known metabolites detected in LC-MS/MS. Alignments of BGCs in the MiBiG database to those identified in a producing strain from this study. Genes are colored according to function. Lines between genes indicate similarity between genes with its color intensity proportional to the BLAST amino acid identity.

### BGCS encoding known antibiotic classes with no metabolites detected - glycopeptide, aminoglycoside and aminocoumarin

In addition to the above-described BGCs whose metabolites were expressed and detected by HRMS/MS, many other BGCs with sequence homology to known antibiotic BGCs were also identified in the sequencing data. However, we were unable to assign metabolite products to these BGCs by LC-HRMS/MS data. A few such BGCs are described below.

Three *Amycolatopsis lurida* strains (GB15-009, GA10-003 and GA10-004) harbored a nearly identical aminocyclitol gene cluster which encoded a homolog of 2-epi-5-epi-valiolone synthase (salQ) responsible for the first step in the biosynthesis of C7N-aminocyclitols (18) (Figure S 9). Aminocyclitols are biosynthesized from sugars through cyclization by a Sugar Phosphate Cyclase (SPC) such as dehydroquinate (DHQ) synthase. The BGCs were highly homologous to cetoniacytone A sharing 70-82% amino acid identity for core biosynthetic genes (19, 20).

In *Amycolatopsis coloradensis* B06-03, an aminocoumarin BGC was detected (Figure S 10). Aminocoumarin antibiotics are biosynthesized from L-tyrosine (21). Tyrosine is activated by an adenylation domain and covalently attached to a peptidyl carrier protein (PCP). A NovH-like cytochorome P450 hydroxylates PCP-bound tyrosine to β-hydroxy tyrosyl-S-PCP. A 3-oxoacyl-acylcarrierprotein (ACP) reductase converts it to a β-keto-tyrosyl intermediate that undergoes cyclization to form 3-amino-4,7-dihydroxycoumarin. In B06-03, downstream core aminocoumarin biosynthesis genes, a type-I polyketide BGC encoding co-enzyme A ligase (CAL) domain specific for 3-amino-5-hydroxybenzoic acid (AHBA), was present as in rubradirin (22) and chaxamycin (23).

A total of seven BGCs similar to previously characterized glycopeptide BGCs were detected (Figure S 11). Glycopeptides are biosynthesized through a multi-modular NRPS assembly line. Glycopeptide BGCs encode additional tailoring enzymes such as P450 monoxygenases and glycosyltransferases that result in amino acid crosslinking and glycosylation respectively, to yield the complex multicyclic antibiotics exemplified by vancomycin. The expected glycopeptides from four glycopeptide-like BGCs (from strains GA10-004, GA10-003, GB8-002, GB15-009) were similar in amino acid composition to ristocetin (24). These four strains primarily contained butylated hydroxytoluene (Bht), dihydroxyphenylglycine (Dhpg) and 4-hydroxyphenylglycine (Hpg) as seen in ristocetin. The predicted glycopeptide from B06-03 is predicted to contain Trp, Hpg and Tyr as seen in complestatin (25).

## Discussion

Soil Actinomycetes hold enormous potential for the discovery of new antibiotics. However, the number of genome-sequenced Actinomycetes in the public domain is still limited, partly due to the cost of long-read next-generation sequencing. In this study, we assessed the capability of the ONT Flongle platform as a low-cost sequencing option to obtain multiple near-complete genomes of Actinomycetes, identify BGCs, and connect them to metabolites through a paired genome-metabolome analysis.

Our sequencing and assembly results showed that up to four near-complete genomes of Actinomycete strains could be sequenced on a single Flongle device. Skipping an optional DNA fragmentation step enabled read lengths up to 80 Kb in each sample. Bead-based size selection further depleted shorter DNA fragments, and enriched sequencing reads in longer sequences (10 Kb+). The long reads enabled contiguous DNA assemblies at lower sequencing depth. The size of the assembly was typical of soil Actinomycetes, suggesting that assemblies represent near-complete genomes of the strains. There were several mismatches in the accuracy comparison analysis in flongle genomes relative to PacBio or high-coverage MinION genomes. However, the contiguity of the genomes was only slightly affected (1-2 contigs verses 6-7 contigs), indicating that important structural information about the genome (e.g., position and organization of genes) can be inferred from these sequences.

A common strategy to obtain contiguous and accurate genome assemblies is through polishing contiguous nanopore assemblies with Illumina reads. One of the significant findings of this study is that BGC predictions and their connection to metabolites was performed without the need for error-correction using Illumina reads. Based on down sampling analysis of public datasets, it was initially hypothesized that low-coverage nanopore-only assemblies could be used to predict and analyze BGCs. Through prospectively sequencing Actinomycetes using Flongle, it was empirically evaluated in the current study.

An interesting observation on BGC analysis was that active site specificity for various BGC classes (NRPS, PKS and CDPS) in the Flongle assemblies were correctly predicted. The active site specificities were used by PRISM to generate possible structures, which were then used to query MS/MS data for potential spectral matches. A spectrum match to a predicted structure indirectly proves that active site specificities were correct, for instance, in the case of cWW. However, we also observed that frameshifts and sequencing errors affected in silico prediction of accurate structures for some BGCs.

The analysis of RiPP BGCs in flongle assemblies was relatively less affected by sequencing. RiPP metabolite prediction is based on short precursor peptide and BGC prediction relies on detecting post-translational modifying enzymes through error-tolerant profile Hidden Markov models. The chance of a mismatch underlying a 100 nucleotide (30-mer core peptide sequence) is low. For example, 19,227 mismatches were detected in total in a flongle assembly relative to PacBio, which corresponds to a chance of less than one mismatch per 100 nt (19,227 mismatches / 7183038 nt genome size × 100 nt = 0.26 per 100 nt). This is evident through accurate prediction of a new lassopepetide BGC and its corresponding experimental mass spectrum in the extract from strain GA6−002.

Similarly, BGCs homologous to previously characterized BGCs for known metabolites can be identified through sequence similarity searches. The consensus accuracy of the assemblies was observed to be 99.5% accurate, which makes a genome sequence suitable for comparison with known BGC sequences or to compute average nucleotide identify with published genomes. We demonstrated this through the rediscovery of BGCs and selected Actinomycetes siderophores.

While our results suggest Flongle is a useful platform for sequencing Actinomycetes, increased consistency in total sequencing output might enable further optimized workflows. For instance, Flongle flow cells were less consistent in the number of starting pores (<60 out of an expected 126 in most cases), which affected total sequencing output and lower than desired coverage for a few samples. A consistent and anticipated number of pores (>100) across experiments would allow for higher sequencing coverage using the same experimental workflow or allow more genomes to be sequenced in an experiment.

While a few BGCs could be connected to the metabolites in the current study, most remained unconnected. Connecting BGCs to metabolites is a multi-factor problem not limited by sequencing accuracy alone. Improvements in experiments and computational algorithms would be needed to circumvent this issue in the future. First, it is highly unlikely that all BGCs will be expressed when strains are grown in only two culture media as tested here; thus, additional media and growth conditions or genetics-free elicitor screens should be used (26–28). Second, only PRISM in silico predicted structures were used. In the future, a more extensive in silico structure generation that addresses ambiguous active site specificities (e.g., two or more possible amino acids at a site in NRPS) could be used. Third, MS/MS data were queried for exact compound spectral matches. Minor differences between predicted and expressed metabolites (e.g., single-site methylation or hydroxylation) would result in a mass shift, and a match would not be possible.

In summary, multiplexed low coverage sequencing of Actinomycetes genomes on Flongle is a promising option for the genome-guided discovery of natural products. Numerous research laboratories house valuable bacterial strain collections (29–34). Limited by the costs of large-scale long read sequencing, genome sequencing of natural product producing bacteria usually occurs on a strain-by-strain basis {Sun, 2021 #295;Li, 2021 #316;Yang, 2021 #317;Braesel, 2018 #326}. The future of natural product research is expected to involve analysis of genomics and metabolomics data using genome mining (e.g. antiSMASH and PRISM) and mass spectrum matching tools (such as molDiscovery integrated with NPAtlas-like databases used in this study). Indeed, such efforts are already taking place on metagenomic data sets (38, 39); while those studies provide vast amounts of data on the natural products-ome, a key limitation is that the data are not connected to archived bacterial strains. It is our hope that low-cost sequencing workflows such as the one described here may allow for access to genome sequencing on a larger scale and/or to a broader community of researchers, especially in resource-limited settings.

## Materials and Methods

### Strain isolation

The twenty sequenced strains were a subset of streptomycin, novobiocin or vancomycin resistant strains from an in-house Actinomycetes strain library housed in the Laboratory of Bioorganic Chemistry, National Institutes of Health. The strains were isolated from soil specimens collected from deserts in Arizona, California and Nevada through standard procedures described in a previous study (35).

### Nanopore sequencing

Each strain was cultivated for 3-7 days in 10 mL of Tryptic Soy broth (BD Diagnostic, catalog no. 211768) with 0.5% (w/v) glycine from frozen glycerol stocks. The cultures were centrifuged at 10,000 x g for 10 minutes and cell pellets were resuspended into 250 μL Tris EDTA (TE) buffer followed by addition of 50 μL of lysosome (100 mg/ml). The mixture was incubated overnight (16 hrs) at 37° C. The next morning 10 μL of RNase A (10 mg/μL) was added to the cell lysate and incubated for an additional 20 min after which 250 μL of proteinase K (400 μg/μL) was added and incubated for 2 hours. DNA was purified from cell lysates using 1:1 v/v phenol-chloroform extraction and the DNA was collected from the upper phase. Genomic DNA was precipitated with 0.7 volume of isopropanol, washed with 80% ethanol, and resuspended into 50 μL TE.

DNA libraries were prepared using Oxford Nanopore Ligation Sequencing Kit (SQK-LSK109) and the native barcoding kit (NBD104) protocol for Flongle with some modifications. A DNA fragmentation step was not performed. 500 ng of genomic DNA was directly processed for DNA end repair with NEBNext Ultra II End repair/dA-tailing Module (New England Biolabs, catalog no. E7546). Barcodes were ligated to the end-repaired DNA and purified with 0.1× or 0.15× beads (Omega Bio-Tek Inc, catalog no. M1378-01), resuspended in a custom buffer (10 mM Tris-Cl pH 8.0, 1 mM EDTA, 0.5 M MgCl_2_ and 5% PEG). A pooled library was prepared by combining 62.5 ng of each barcoded DNA. Nanopore adapters were ligated to the pooled library followed by library loading and sequencing according to the manufacturer’s instructions.

### Data-dependent untargeted LC-MS/MS

Each strain was cultivated in deep well plates containing 400 μL of ISP1 (BD Diagnostic, catalog no 276910) or R2A media (Teknova, catalog no. R0005). The cultures were incubated at 30° C with shaking at 200 rpm for one week before extraction with an equal volume of ethyl acetate followed by extraction with *n*-butanol. Uninoculated media were used as blanks / negative control, and any metabolite observed in a blank run was excluded from interpretation. The LC-MS/MS data was collected using an Agilent 1290 Infinity II UPLC system equipped with an Agilent 6545 qTOF mass spectrometer. Samples were chromatographed on a Agilent Eclipse Plus C18 2.1×50mm column (3 μL injections) using a gradient of 99% A (0.1% formic acid in water) to 95% B (acetonitrile) at a flow rate of 0.5mL/min over 10min. MS/MS fragmentation was carried out in auto mode with collision energies of 10, 20 and 40 KeV excluding precursor ions in the range of 40-180 *m/z* and abundance below 7,000 counts.

### Data analysis

#### Genomics

Primary genomic data analysis was conducted by basecalling with guppy (version 4.2.2, model dna_r9.4.1_450bps_hac.cfg), demultiplexing with Qcat (version 1.0.6) and assembling with Canu (version 2.0) (40). Canu assemblies were constructed with an expected genome size of 8 Mb, minimum read length threshold of 1 Kb, minimum coverage of 2, and high Mhap error correction sensitivity (40). The genome assemblies were polished by aligning reads to the assembly and calling consensus with Racon and Medeka (41). Genes and secondary metabolite gene clusters were predicted using the programs antiSMASH (version 5) and PRISM (version 4.4.5) (11).

### Assessing effect of sequencing coverage on BGC detection

FastQ reads for previously sequenced Actinomycete genomes were downloaded from the European Nucleotide Archive, dowsampled to an estimated coverage of 60×, 30×, 15× and 7× using seqtk (assuming an 8 Mb genome as in prospective sequencing) (Table S 1). Seqtk allows random subsampling of reads. Reads were subsampled to desired coverage according to the following estimation: number of reads for Q coverage = (Q/original coverage) × original number of reads. The downsampled FastQ files were assembled with Canu, polished with Medeka and BGCs predicted using antiSMASH. The number of mismatches in each assembly relative to original coverage was calculated using Quast (42). For consistency with data from this study (presented in Figure 3), the coverage presented is aligned coverage, taking into account the size of the final assembly and not the expected size (i.e. 8 Mb). The mapped coverage was extracted from Canu assembler tig information files.

### BGC comparison between strains

The antiSMASH-predicted BGC sequences were extracted from each strain’s genome assemblies and aligned in all possible strain pair combinations using minimap2, allowing for 5% sequence divergence (43). If a BGC from strain-1 did not align to any of the BGCs in strain-2, it was considered absent in strain-2.

### Homologous BGCs of previously characterized metabolites

Homologous BGCs of previously characterized metabolites were obtained through the ‘known cluster blast’ module of antiSMASH. The output of the program contains gene-wise blast hits for each BGC in the genome to BGCs in MiBiG (4). To identify the best hit in MiBiG database, output was first filtered to obtain MiBiG BGCs that share the largest number of genes, highest mean percent identity and highest mean coverage with genes in the query BGC. The results were subsequently filtered to retain only BGCs where the ratio of lengths between query and MiBiG BGCs was between 0.7 to 1.1.

#### LC MS/MS Analysis

Analysis of LC MS/MS data was conducted by conversion of the vendor.d format to mzXML files using the GNPS conversion utility. These mzXML files were subsequently used for all analyses.

### Spectrum matching - known or unknown structures

Spectrum matches for known metabolites using the Natural Product Atlas or unknown metabolites (PRISM predicted structures) were identified by using molDiscovery (3, 12). molDiscovery computes theoretical MS/MS spectra of compounds in the database, identifies spectrum matches at user-defined mass-tolerance, and subsequently calculates statistical significance by matching the spectrum against a decoy database. In this analysis, mass tolerance of 20 ppm, p-value less than e^−10^ and FDR less than 1% were considered.

### RiPPs

Spectrum matches for RiPPs were detected using metaminer (14). Given a list of short peptides, metaminer constructs possible RiPP products based on knowledge of post-translational modifications within RiPPs. It then predicts an MS/MS spectrum for each predicted RiPP product and conducts a search of experimentally collected MS/MS spectra for potential matches. All open-reading-frames (ORFs) shorter than 600 nt (200 amino acids) located within RiPP BGCs predicted by antiSMASH or PRISM were used for this analysis.

Integration of the mass spectrometry and genomic sequences was achieved through R scripts and several packages including MSnbase (44) and Open Babel (45).

### Data availability

Raw sequencing data are available under NCBI Project accession no: PRJNA752621. Genome assemblies and additional data are available at Figshare (https://figshare.com/articles/dataset/_/15094044). The MS/MS spectra have been uploaded to GNPS with accession number: MSV000087950. Scripts used in data analysis and preparation of figures are available at https://github.com/rajwanir/flongle_actinomycetes_paper.

## Acknowledgements

This work was supported by the NIH Intramural Research Program (NIDDK) and utilized the computational resources of the NIH HPC Biowulf cluster (http://hpc.nih.gov).

